# Escalation and constraints of antagonistic armaments in water striders

**DOI:** 10.1101/430322

**Authors:** Antonin Jean Johan Crumière, David Armisén, Aïdamalia Vargas-Lowman, Martha Kubarakos, Felipe Ferraz Figueiredo Moreira, Abderrahman Khila

**Affiliations:** Institut de Génomique Fonctionnelle de Lyon, Université de Lyon, Université Claude Bernard Lyon 1, CNRS UMR 5242, Ecole Normale Supérieure de Lyon, 46, allée d’Italie, 69364 Lyon Cedex 07, France; Section for Ecology and Evolution, Department of Biology, University of Copenhagen, Universitetsparken 15, 2100 Copenhagen, Denmark; Department of Pharmaceutical Sciences, University of Toronto, Toronto, Ontario M5S 3M2, Canada; Laboratório de Biodiversidade Entomológica, Instituto Oswaldo Cruz, Fundação Oswaldo Cruz, Rio de Janeiro 21040-360, Brazil

**Keywords:** Sexual conflict, antagonistic armaments, coevolution, water striders, sexual dimorphism

## Abstract

Sexual conflict may result in the escalating coevolution of sexually antagonistic traits. However, our understanding of the evolutionary dynamics of antagonistic traits and their role in association with sex-specific escalation remains limited. Here we study sexually antagonistic coevolution in a genus of water striders called *Rhagovelia*. We identified a set of male grasping traits and female anti-grasping traits used during pre-mating struggles and show that natural variation of these traits is associated with variation in mating performance in the direction expected for antagonistic co-evolution. Phylogenetic mapping detected signals of escalation of these sexually antagonistic traits suggesting an ongoing arms race. Moreover, their escalation appears to be constrained by a trade-off with dispersal through flight in both sexes. Altogether our results highlight how sexual interactions may have shaped sex-specific antagonistic traits and how constraints imposed by natural selection may have influenced their evolution.

## Introduction

The evolutionary interests of males and females in reproductive interactions often diverge, leading to the coevolution of sexually antagonistic traits that are favoured in one sex at a fitness cost to the other (1). The evolution of these dimorphic traits can, in turn, cause antagonistic selection and episodes of escalation of armaments between the sexes (2, 3). Conflict resolution, through the evolution of sex biased gene expression and sexual dimorphism, is expected to drive divergence of the sexes sometimes resulting in striking cases of sexual dimorphism (4, 5). In the specific case of sexual conflict over mating rate, the evolution of antagonistic armaments, such as grasping traits in males, can be matched by the evolution of counter-adaptations as anti-grasping traits in females (6, 7). These sex-specific armaments can lead to episodes of escalation and ultimately arms race (3, 8).

Antagonistic co-evolution of the sexes has attracted much attention and many studies focused on sex chromosome evolution (3, 9), sex biased gene expression (10–12), and on the dynamics of morphological co-evolution of sexually antagonistic traits (2, 13, 14). However, our understanding of the function of sexually antagonistic traits during interactions between the sexes, their impact on the outcome of sexual interactions, and the factors that escalate or constrain the expression of these traits remain unclear. Waters striders have been widely used as a model to study sexual dimorphism and antagonistic co-evolution (7, 8, 15). Sexual dimorphism is particularly dramatic in the tropical genus *Rhagovelia* (Insecta, Gerromorpha, Heteroptera) (16–18), where both sexes often bear morphological traits reminiscent to those found in species with strong sexual conflict (1, 16–18). However, the process of sexual selection driving sexual dimorphism in *Rhagovelia* is unknown (1, 19). Here, through morphological and behavioral quantifications, we identified a large set of potentially sexually antagonistic armaments in the *Rhagovelia* genus and tested their role during sexual interactions in a representative species called *Rhagovelia antilleana* (16–18). We then tested how variation in sexually antagonistic traits affects male and female success during sexual interactions. We show that this variation is associated with trade offs with flight capability and egg storage. Finally, we used phylogenetic reconstruction of male and female secondary sexual traits to understand how antagonistic coevolution and diversification may have shaped the divergence of the sexes in this lineage and assess the extent of evolutionary arms race.

## Results

### Sexual dimorphism in *Rhagovelia*

Sexual dimorphism in the water strider genus *Rhagovelia* ranges from subtle to spectacular differences in morphology between the sexes (Figure 1A-D’). *Rhagovelia* populations contain both winged and wingless morphs (17, 18). In both male morphs, the foreleg tarsi are equipped with prominent, hook-like, combs that are absent in the females (Figure 1 E and E’). The rearlegs of males possess several rows of spikes of different sizes along the trochanter, femur, and tibia (Figure 1F, F’ - G, G’). In addition, the femurs of the rearlegs are very thick compared to those of the females (Figure 1F, F’). Staining for actin fibres revealed that the width of the femur is entirely occupied by muscles (Figure S1), suggesting that thicker femurs have the potential to apply stronger grip force (Figure 2D-F). In females, winged morphs possess a prominent spike-like projection of the pronotun (Figure 1 H’) that is not found in wingless females (Figure 1C’) or winged males (Figure 1H). Wingless females have a narrow-shaped abdomen (Figure 2B) that can be even more pronounced in some species such as *Rhagovelia obesa* (Figure 2C). We have never observed winged females with narrow abdomen or wingless females with pronotum projection indicating that these two traits are mutually exclusive (Figure 2A-B). This observation suggests that female morph-specific strategies to reduce mating frequency have evolved in *Rhagovelia* females.

**Figure 1:**
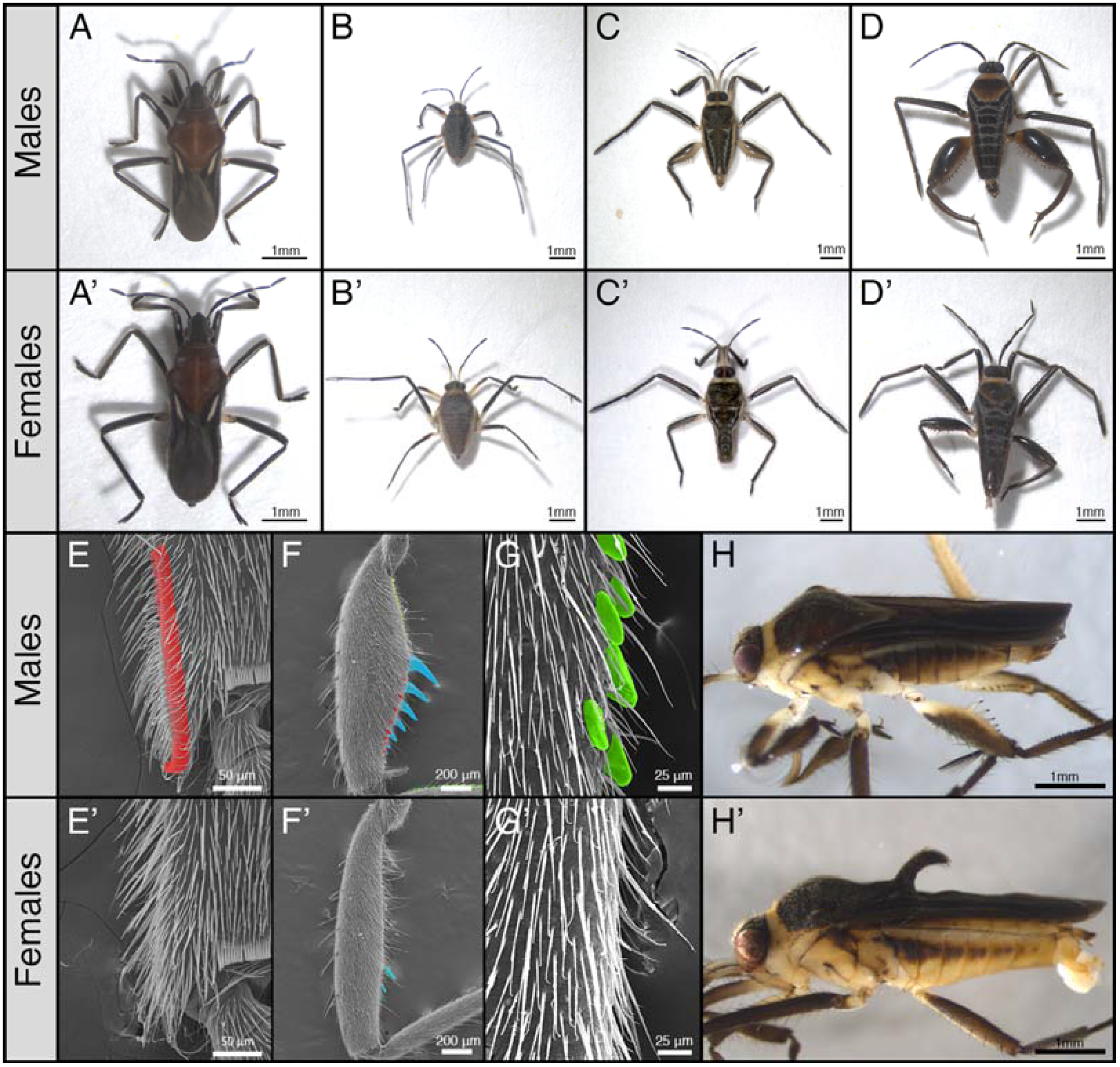
Sexual dimorphism and secondary sexual traits in the genus *Rhagovelia*. The outgroup *Oiovelia cunucunumana* (A, A’) does not show strong dimorphism. In the genus *Rhagovelia*, sexual dimorphism can be subtle as in *Rhagovelia plumbea* (B, B’) or more extreme as in *Rhagovelia antilleana* (C, C’) or *Rhagovelia* sp.2 (D, D’). This dimorphism affects secondary sexual traits such as the sex combs (in red) present in male fore-legs (E) and absent in females (E’); the presence of spikes (in blue, yellow and red) and the larger femur of the rear-leg femur in males (F) compared to females that only have some small femur spikes (F’); and the presence of spikes (in green) along the rear-leg tibia in males (G) that are also absent in females (G’). Females possess secondary sexual traits such as a narrow abdomen (C’) and a pronotum projection (H’) that are absent in males (C, H).

### Mating behaviour in *Rhagovelia antilleana*

Pre-mating interactions in *R. antilleana* include vigorous struggles between males and females characteristic of sexual conflict over mating rate (1, 8, 20). Behavioural quantification revealed that males attack females preferably on the side by grasping female’s mid- and rearlegs (328 out of 461 attacks, Table 1). We did not detect a significant male preference for winged or wingless morphs (Figure S2A; n=36 winged females and n=36 wingless females; Wilcoxon test: p=0.94 n.s.), however we observed that wingless females were mainly attacked from the side (63.2%) and the back (29.6%), while winged females were more likely to be attacked from the side (78.5%) possibly due to the presence of the pronotum projection (Table 1). When mounted, males use the sex combs to clasp the pronotum of the female and tighten the grip on her legs using their modified rearlegs (Supplementary video 1). Females shake their body vigorously and perform repeated somersaults, which frequently result in rejecting the male (Supplementary video 2, Table 1). The struggles are vigorous and their duration is typically 0.68 +/-0.99 seconds in average, but can vary between fractions of a second (<0.15 seconds) to several seconds (9.75 seconds) (Figure S2B). These pre-mating struggles are indicative of sexual conflict over mating rate and the sex-specific modifications we observed might be the result of antagonistic interaction between the sexes in this species.

**Table 1:**
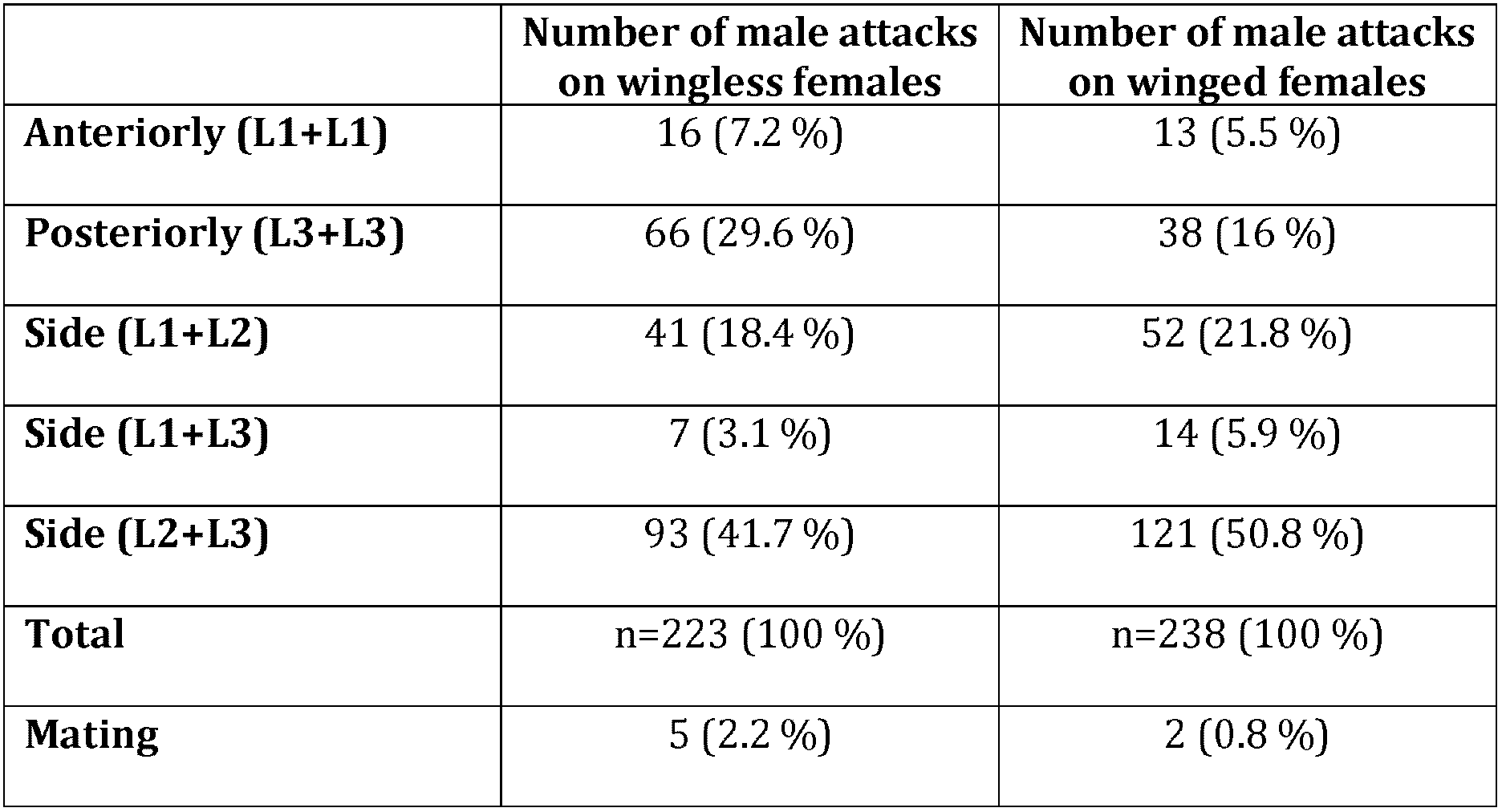
Quantification of preferential way of attack used by males on different female morphs. Males preferentially attack females on the side rather than anteriorly or posteriorly. This tendency increases when males try to mate with a winged female. A total of 223 and 238 interactions with wingless and winged females respectively were observed during 18 trials.

### Effect of natural variation in male and female antagonistic traits on mating rate

Artificially generated variation in sexually antagonistic traits is known to results in variation in mating success (20). Here, we wanted to test whether natural variation in male persistence and female resistance traits is also associated with variation in mating success. In *R. antilleana*, only winged females have the pronotum projection (Figure 2A) and only wingless females have narrow abdomen (Figure 2B-C), whereas in males there is considerable natural variation in the degree of elaboration of the rear-legs (Figure 2D-F). To determine the extent to which this variation affects individual performance during pre-mating struggles, we used a tournament design (Figure S3) (21) composed of 10 replicates with 8 males and 8 females each for a total of 80 males and 80 females. This experiment separated males and females based on the outcome of premating struggles into successful, intermediate, and unsuccessful individuals (Figure S3, Figure 3). Successful males are those who copulate after a premating struggle and successful females are those who reject males after pre-mating struggle (Figure S3). We found that successful males have significantly more rear-leg tibia spikes and thicker rear-leg femurs compared to unsuccessful males (respectively Wilcoxon test: p = 0.045*; Student t-test: p = 0.02*; Figure 3A and B), however body size, number of spikes on rear-leg femur, and the length of rear-leg tibia and femur did not significantly differ between the two groups (Figure 3 C-F).

**Figure 2:**
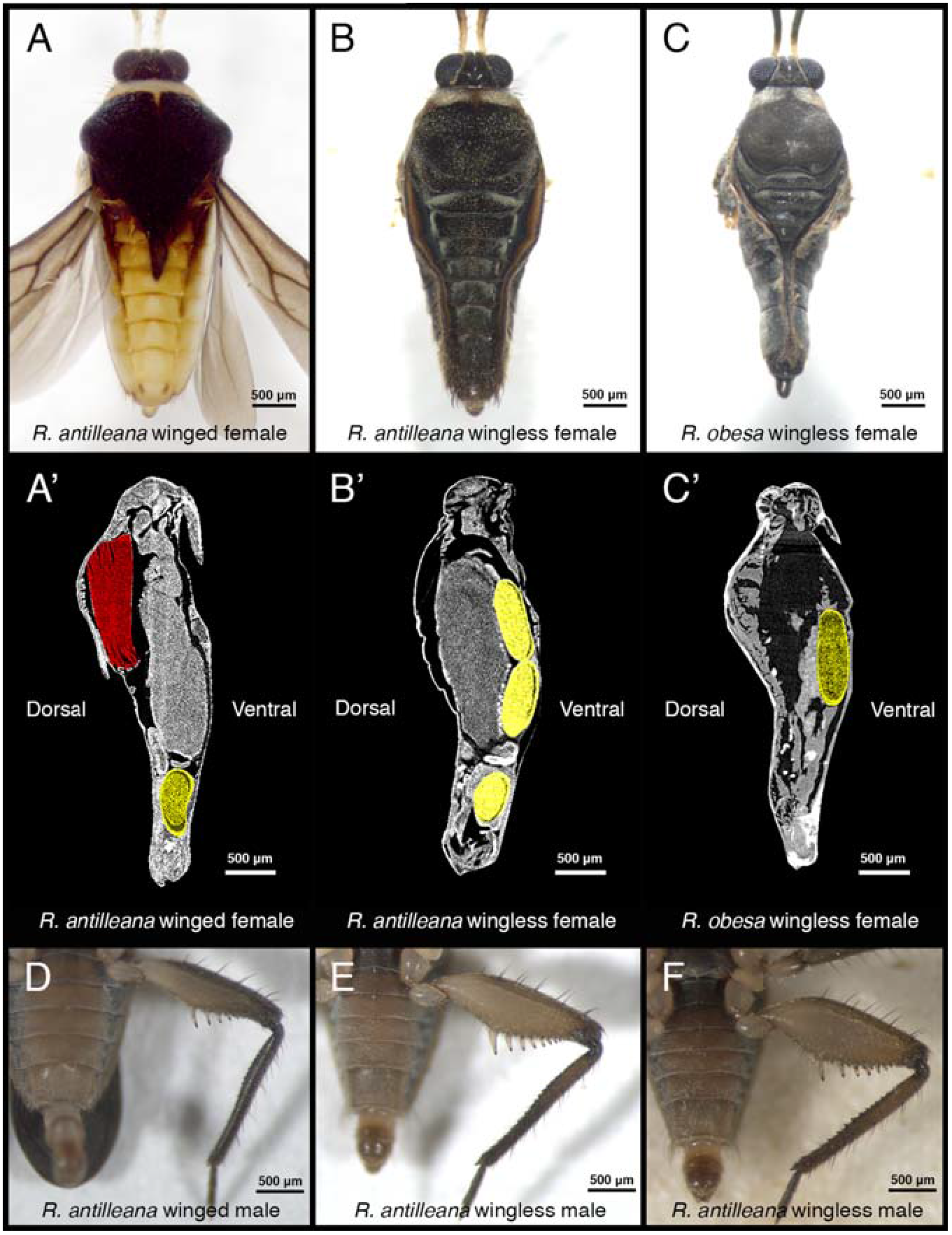
Variation in secondary sexual traits in *Rhagovelia*. Winged *R. antilleana* females (A) have a large abdomen compared to wingless females (B). The presence of wing muscles (A’, in red) in the thorax constrains the eggs to be located in the abdomen (in yellow) while in wingless females (B’) the space in the thorax is free from wing muscles and can be occupied by eggs, thus allowing the development of a narrow abdomen. In species such as *R. obesa* (C), the narrow abdomen is even more pronounced and eggs are located in the thorax (C’), thus confirming our observation. There is also variability in the rear-leg in males, especially in the size of the femur. Winged males (D), have thin femurs while other males have intermediate (E) or large femurs (F).

This result highlights the importance of the rear-legs for increasing male mating frequency and therefore male fitness (1). In females, our data indicated that individuals with a pronotum projection were significantly more efficient in rejecting males than females with narrow abdomen (Cochran-Mantel-Haenszel chi-squared test: 23,45; df = 2; p-val: 8.089e-06***; Figure 3G). To further test the validity of this conclusion, we conducted an experiment where the performance of 30 wingless females (narrow abdomen and absence of pronotum projection) was compared to that of 18 winged females (wide abdomen) from which we amputated the pronotum projection. We found that winged females with amputated pronotum projection were less efficient at rejecting males, which resulted in higher mating frequently (Figure 3G-H). This result indicates that in the absence of the pronotum projection, narrow abdomen increases female’s ability to resist harassing males. Therefore, alternative morph-specific strategies evolved in the females possibly under selection by male harassment. Altogether, these experiments demonstrate that natural variation in sexually antagonistic traits is associated with variation in the ability of both sexes to control mating rate.

**Figure 3:**
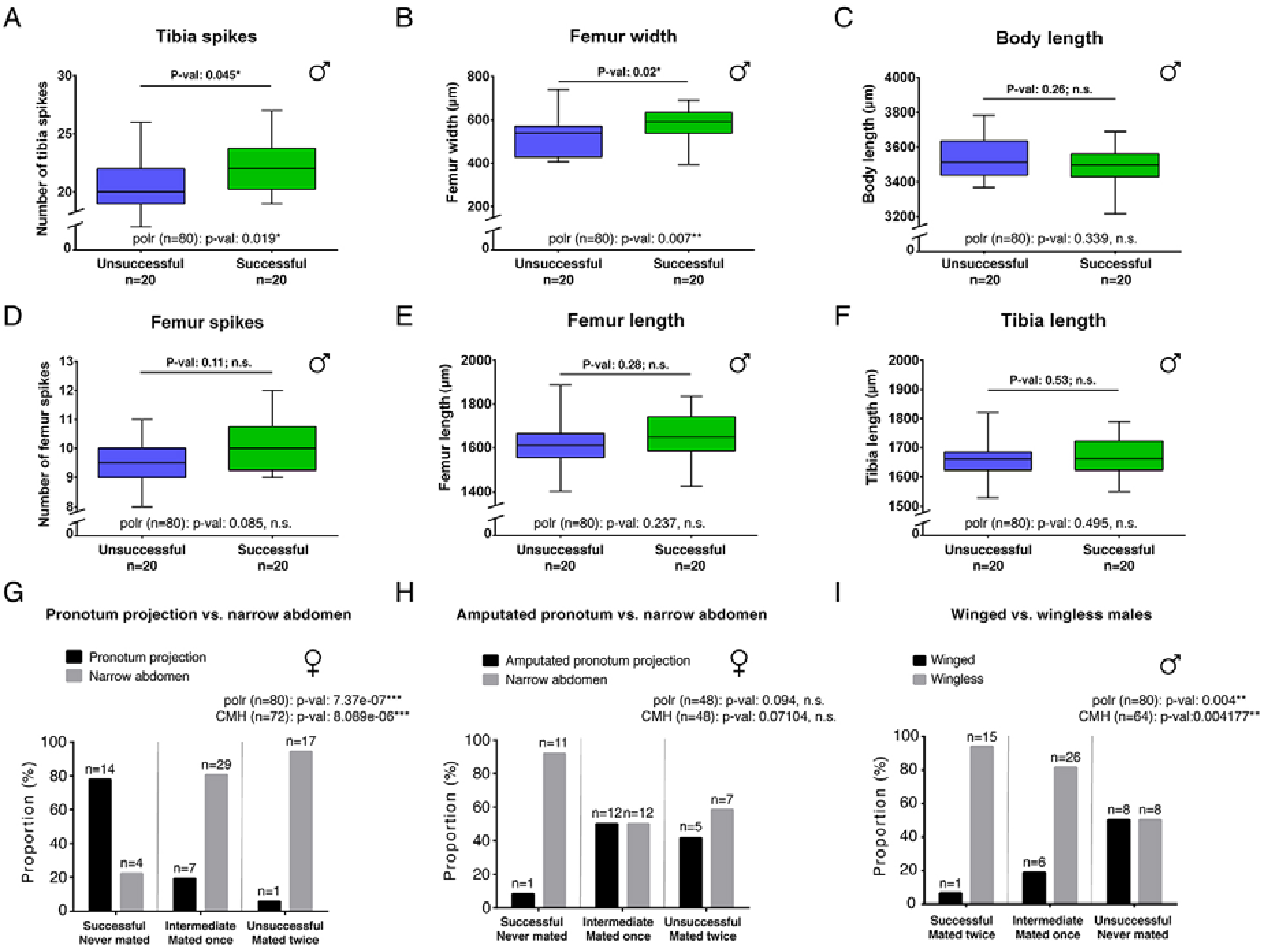
Morphological comparisons and effect on mating performance. Successful males have significantly higher number of tibia spikes (A) and larger femur (B) compared to unsuccessful males. There are no significant differences between the two classes of males for body length (C), number of femur spikes (D), femur length (E) and tibia length (F). Student t-tests were performed for all comparisons except for the number of femur spikes (Wilcoxon test). (G, H and I) show the distribution of the different morphs ecotype in the different classes of performance with the associated results of Cochran-Mantel-Haenszel Chi-square test. Significant p-values indicate association between the morph ecotype and the performance. Females with pronotum projection are more efficient than females with narrow abdomen in rejecting males (G). Females with amputated pronotum projection lost their ability to efficiently reject males (H). Wingless males are more efficient than winged males in mating (I). A *proportional odds logistic regression* analysis (polr) for the different phenotypic variables is indicated in all graphs. Significant p-values indicate association between the phenotypic trait measured and success in mating efficiency for males and male rejection for females. *n* indicates sample size.

### Armament escalation is constrained by flight capability in *Rhagovelia*

Because natural variation in antagonistic traits can result is variation in the performance of males and females during pre-mating struggles, we investigated which factor might influence this variation. Many studies described the presence of trade-offs between flight capability and certain life history traits such as fertility in a number of insects (22–24). We therefore tested whether wing polymorphism in *Rhagovelia antilleana* affects variation in male’s and female’s sexually antagonistic traits. The pronotum projection, a sexually antagonistic trait that increases the ability of females to resist male harassment, only develops in winged female morphs (Figure 1H’). The development of the wings is accompanied by the development of flight muscles, which fill most of the space in the thorax (Figure 2A’) (25–27). These winged females position their developing eggs mostly in the abdomen (Figure 2A’). In wingless females of both *Rhagovelia antilleana* and *Rhagovelia obesa*, we found that the absence of flight muscles provides a space in the thorax that is occupied by the developing eggs (Figure 2B’, C’). Furthermore, we found that wingless females contained a significantly higher number of developing eggs than winged females, consistent with the known trade off between flight and fertility (respectively 4.15 mean ± 2.96 SD; n=20 and 1.35 mean 2.35 ± SD; n=20; Student t-test:p-val 0.002094**; Figure 2A’-C’). These data suggest that the presence of wings favours the development of the pronotum projection and constrains the narrowing of the abdomen (Figure S4). These constraints may have driven the evolution of alternative sex-specific and morph-specific strategies to escape fitness costs due to frequent mating in females. In males, we observed that winged morphs typically have the thinnest rear-leg femurs (Figure 2D) and were under-represented in the successful group of our tournament set-up, contrary to wingless morphs that have the largest rear-leg femurs (Cochran-Mantel-Haenszel chi-squared test: 10,957; degree of freedom: 2; p-val: 0,004177**; Figure 3I). These observations highlight a trade-off between dispersal and mating success also in males. Altogether, these results suggest that some life history traits such as fertility and dispersal constrain the escalation of sexually antagonistic armaments in both sexes (28, 29).

### Signs of evolutionary arms race between the sexes in the *Rhagovelia* genus

Coevolution of the sexes due to antagonistic interactions could lead to escalation and arms race that deeply shape the evolutionary trajectory of lineages in nature (2, 6, 8, 13, 14). We therefore tested the evolution of armament and presence of arms race in the *Rhagovelia* genus by analysing phylogenetic patterns of correlation of male and female phenotypic complexity in terms of secondary sexually antagonistic traits (2, 13, 14). We generated a matrix of these traits in both males and females in a total of thirteen species including nine *Rhagovelia* and four closely related outgroups (Table S1 and S2). We mapped both male and female traits on a phylogeny to determine the patterns of phenotypic complexity between the sexes (Figure 4, Figure S5, Figure S6). Our reconstruction showed a strong phylogenetic signal where males of the *Rhagovelia* genus evolved an increasing number of secondary sexual traits (Figure 4). This pattern indicates substantial escalation of conflict in males (2, 6, 13, 14). In females, we detected the evolution of anti-grasping traits in a clade where males are the most armed (Figure 4), suggesting female counter-adaptation to male escalation (2, 6, 13, 14). Altogether, these patterns of armament of the two sexes suggest an on-going arms race in the *Rhagovelia* genus.

**Figure 4:**
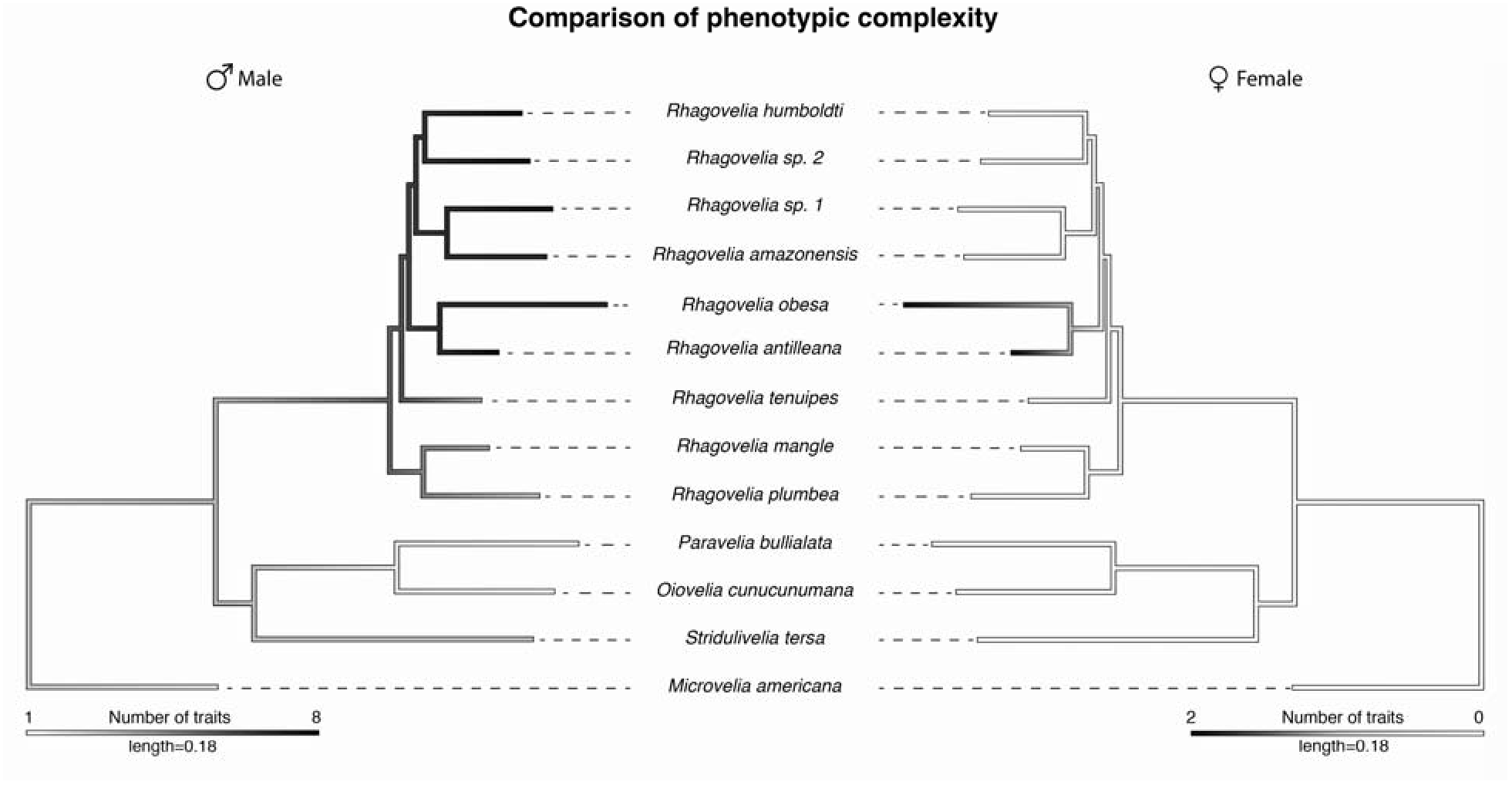
Comparison of male and female phenotypic complexity. The phylogeny of our samples shows a higher complexity of male phenotypes, in terms of number of secondary sexual traits, compared to outgroups (left phylogeny). This complexity is followed by higher complexity in females in two species *R. antilleana* and R. *obesa*, the ones that possess narrow abdomen and pronotum projection, suggesting an ongoing arms race.

## Discussion

We have shown that sexual conflict over mating rate generates sexual dimorphism via the development of secondary sexual traits in the *Rhagovelia* genus. Males have evolved a set of persistence grasping traits aimed at increasing mating, whereas females evolved a set of anti-grasping traits aimed at decreasing mating. Natural variation in these antagonistic traits directly impacts mating frequency and therefore influences the fitness of both sexes. Among male grasping traits are the sex combs known in other insects (30–34). An additional male set of grasping traits consists of elaborations in the rearlegs, which are used as clasps that neutralize female’s legs (Supplementary video 1). Our analysis revealed that higher number of spines on the tibia and larger femur are the two main structures associated with male mating success. Females on the other hand, possess morph-specific modifications consisting of a narrow abdomen in wingless morphs and a pronotum projection in winged morphs. Both of these traits increase the chances for the female to reject harassing males. The projection forms a barrier and increases the distance between the abdomen of the male and the female, likely making it easier for the female to resist. Our experiments also showed that females with narrow abdomen are better at rejecting males that females to which we removed the pronotum projection and that have a wide abdomen. Because males use their rearlegs as clamps on the legs of the female, it is possible that the narrowing of the abdomen allows the female to slide out of this grip. Altogether our observations indicate a role of these sex-specific phenotypes in male persistence and female resistance (2, 6).

Our results also indicated that dispersal, manifested by the presence of wings (16, 35, 36), constrains the degree of expression of sexually antagonistic traits in both sexes. This constraint reduces elaboration that in turn reduces the ability of the sexes to control mating frequency. Therefore, there is a tradeoff between dispersal and the expression of sexually antagonistic traits in *Rhagovelia*. Interestingly in the case of winged females, wing development allows the extension of the pronotum, which is an efficient female resistance trait. This suggests that the development of this structure might be contingent on the activation of the wing development program, although this hypothesis remains to be experimentally tested. How the decision to develop into winged or wingless morphs and whether sexual conflict affects this decision in *Rhagovelia* is unknown. *Rhagovelia* is a tropical genus and populations occur in large groups that can reach hundreds of individuals, suggesting that male harassment is frequent (37). It is possible that under these conditions, winged morphs are favored to allow for dispersal and occupation of niches with lower population densities. However, winged morphs may not be favored in other contexts because of the trade-offs we observed between the presence of wings, the number of eggs produced by females, and the low success in pre-mating interactions we observed for males. It is possible that trade-offs, between mating frequency, dispersal, and fertility have shaped phenotypic evolution in this group of insects.

Finally, antagonistic sexual interactions and constraints have driven evolutionary escalation in the *Rhagovelia* genus (2, 6). Our analysis has shown that, across this genus, males have evolved an arsenal of traits across different species while females only evolved two morph-specific armaments (i.e. narrow abdomen and pronotum projection) in only in two species from a single clade in our sample. These are *Rhagovelia antilleana* and *Rhagovelia obesa* both belonging to the *collaris* complex (18). The presence of modified females in this group, with the pronotum projection in winged females and narrow abdomen in wingless females (18), suggests that sexual antagonism had strongly impacted the evolution of the genus. In addition to morphological modifications, the females have evolved complex behavior among their resistance arsenal through shaking and vigorous somersaults. It would be interesting for future studies to investigate the difference in female resistance behavior across species and assess to which extant females rely on behavior relative to morphological modifications.

## Conclusion

Our approach has linked behavioral quantifications, morphological observations, mating performance assessment and phylogenetic analysis for a better understanding of morphological evolution and diversification. The results we obtained through this study highlight how sexual interactions and natural selection can enhance and constraint antagonistic coevolution between males and females and participated to phenotypic diversification of closely related species.

## Methods

### Insect sampling and culture

Species were collected during fieldwork in the locations indicated in Table S1. Lab populations were established for *Rhagovelia antilleana*. They were kept in water tanks at 25°C, 50% humidity, 14 hours of daylight and fed daily on crickets. Styrofoam floaters were provided for adult female egg laying. Adults and nymphs were raised in independent tanks to decrease nymph cannibalism.

### Imaging

Picture acquisition and observation of secondary sexual traits were performed using a SteREO Discovery V12 (Zeiss), a Stereomicroscope M205 FA (Leica) and using Scanning Electron Microscopy at Centre Technologique des Microstructures (UCBL).

### Muscle staining

Male rear-leg femurs were dissected and opened using forceps and scalpel to remove the cuticle. Femurs were fixed with Formaldehyde 4% in PTW 1% during 20 minutes. Femurs were then washed in 5 successive 10-minute baths of PTW 1%. The muscles were marked with 1/1000 Phalloidine 488 (Invitrogen) for 1 hour followed by 5 successive 10-minute baths in PTW 1%. The femur were mounted in 50% glycerol + Dapi.

### Analyse of male behaviour during pre-mating struggles

We observed 36 non-virgin couples composed of one male and one female (18 wingless and 18 winged females) using the Phantom Miro M310 Digital High Speed Camera and PCC Software (Vision research, Ametek) for video acquisition. We recorded 223 interactions between males and the wingless females and 238 between males and the winged females. We then analysed how males attack the female and classify the different types of attack into “anteriorly” if males attack from the front and block the female T1-legs (fore-legs), “posteriorly” if males attack from the back and block the female T3-legs (rear-legs), and from the side if the males block female T1- and T2-legs (mid-legs), T1- and T3-legs or T2- and T3 legs.

### Determining male preference for female morphs

To determine male preference over types of female morphs we have put 2 non-virgin males with non-virgin females (one winged and one wingless) into a common plastic container with water and recorded the number of interactions with males for each female morphs during 20 minutes. We repeated this procedure 18 times for a total of 36 winged and 36 wingless females.

### Recording duration of interactions related to female morphs

To determine if the pronotum projection in females influence the duration of interactions, we recorded the time of the interactions for 36 non-virgin couples composed of one male and one female (18 wingless and 18 winged females) during 20 minutes each using a stopwatch with centisecond option. Interactions shorter than 0,15 seconds were too fast to take not and were not take into account in the analysis of data. A total of 604 and 489 interactions were recorded for respectively winged and wingless females.

### Quantification of trait functions

Non-virgin males and females were isolated into two different buckets, one for each sex, during 5 days to increase the motivation to mate. We set-up 10 tournaments between 8 males and 8 females (80 males and 80 females in total) to split them in three different categories: successful, intermediate and unsuccessful, depending the ability in copulating for males and the ability in rejecting male for females. Both males and females were introduced at the same time in a common bucket of water. During the first round, for each mating we removed the couple and interrupted the mating by separating the male and the female in order to preserve the motivation to mate for the second round. The male was classified as successful and the female as unsuccessful. Inversely, the remaining males that did not succeed to mate were classified as unsuccessful in mating and remaining females as successful in rejecting. The first round goes on until we obtained 4 successful males and 4 unsuccessful females (Figure S3). The different rounds of this experiment occurred for few minutes to several hours depending on the activity of the bugs. For the second round, we used two different buckets. In the first bucket, we introduced, at the same time, the 4 successful males with the 4 successful females, and in the second bucket we introduced, at the same time, the 4 unsuccessful males with the 4 unsuccessful females. At each mating, we removed the couple, as in the first round. The second round goes on until we obtained 2 successful males and 2 unsuccessful females per buckets. At the end, we were able to obtain the following rank for both males and females: successful for males that have mated twice and females that have never mated, intermediate for individuals that mated once, and unsuccessful for males that have never mated and for females that mated twice (Figure S3). Then, using SteREO Discovery V12 (Zeiss) with ZEN 2011 software (Zeiss) we took pictures of each individual to quantify the individual phenotypes. We measured the length and the width of the rear-leg femur, the length of the rear-leg tibia and we counted the number of spikes on the rear-leg femur and the number of spikes on the rear-leg tibia. We also recorded the presence of a pronotum projection, narrow abdomen and body length in females. To further test the role of the pronotum projection, we manually removed the pronotum projection from 18 winged females using forceps and kept these females in lab condition during one week to allow them to recover. Then we set-up 6 new tournaments to compared the ability of these 18 winged females in rejecting males against 30 wingless females. All data are available in the Dryad Digital Repository: *Will be provided before publication*.

### Statistical analysis

Cochran-Mantel-Haenszel chi-squared tests (Figure 3 G-I) were performed on data of the tournament using only the replicates where both morph types were present in order to compare performance (i.e. winged vs. wingless; 9 replicates (72 females) for pronotum projection vs. narrow abdomen; 6 replicates (48 females) for amputated pronotum vs. narrow abdomen; 8 replicates (64 males) for winged vs. wingless morphs). Proportional odds logistic regression (MASS package) (38) were performed on data of the tournament using all individuals in order to analyse phenotypes related to mating performance. Shapiro tests, and Student t-tests and Wilcoxon tests for analysis of male preference, duration of interactions, comparison of trait values for successful and unsuccessful males, depending whether variables followed normal distributions, were performed using RStudio version 1.0.153. R Script used in this study is available in the Dryad Digital Repository: *Will be provided before publication (or upon request)*.

### Phylogenetic reconstruction

Sequences were retrieved from in house transcriptome databases and from (39) for the following markers: *12S RNA; 16S RNA; 18S* RNA; *28S RNA; Cytochrome Oxydase subunit I (COI); Cytochrome Oxydase subunit II (COII); Cytochrome Oxydase subunit III (COIII); Cytochrome b (cyt b); NADH-ubiquinone oxidoreductase chain 1 (ND1); Ultrabithorax (Ubx); Sex combs reduced (Scr); Gamma interferon inducible thiol reductase (gilt); Antennapedia (Antp); Distal-less (dll); Doublesex (dsx)*. All these markers were submitted to GenBank and their accession numbers can be found in the Dryad Digital Repository: *Will be provided before publication (or upon request)*. Phylogenetic reconstruction (Figure S5) was performed using MrBayes version 3.2.6 and PhyML version 3.0 in Geneious 7.1.9 as described in (39). Concatenation of sequence alignments and phylogenetic tree in Newick format are also available in the Dryad Digital Repository: *Will be provided before publication (or upon request)*.

### Assessment of armament escalation

We created a matrix of presence/absence of secondary sexual traits for both males and females (Table S2). Based on our observation, we found eight traits in males; the sex combs in the forelegs, the different rows of spines (up to 5 rows) and the shape of the different segments of rear-legs. In females, these traits include the pronotum projection and the narrow abdomen shape. Then, we mapped the sexually antagonistic traits individually on our phylogeny to reconstruct the ancestral state of each of them (Figure S6). Finally, we performed a reconstruction using the combination of all traits, in males and females separately, to determine the pattern of phylogenetic complexity in this sample. Reconstruction of ancestral state was performed using Mesquite version 3.2 (40) and mapping of phenotypic complexity was performed using contMap (package phytools, (41)) in RStudio version 1.0.153.

### Observation and imaging of egg position and egg counting

We used CT Scan to observe egg position without disrupting internal morphology of our samples. We fixed females of *R. antilleana* (n= 5 winged; n=5 wingless) and *R. obesa* (n=5 wingless) during 2 days using Bouin solution (MM France). Then, samples were washed twice with PBS 1X and were emerged 4 days in 0,3% phosphotungstic acid + 70% ethanol + 1% Tween20 solution to improve the contrast during microtomography acquisition (42). Specimens were scanned using Phoenix Nanotom S (General Electrics) using the following parameters: 30 kV tensions, 200 μA intensity, 2400 images with time exposure of 2000 ms and a 2.5 μM voxel size. Three-dimensional images were reconstructed with the software attached to the machine (data rec) and then visualized with VGStudioMax. Longitudinal pictures were acquired using this same software to assess the presence of wing muscles and the position of eggs. Further egg counting was performed after dissection of 20 winged and 20 wingless *R. antilleana* females.

### Two-dimensional morphometric analyses of body shape

Pictures of adult females of *R. antilleana* (n= 10 winged; n=10 wingless) and R. *obesa* (n=9 wingless) were acquired using SteREO Discovery V12 (Zeiss). The body outline of each individual was extracted using Adobe Illustrator CC 2017 (Adobe). The analysis of outlines of the different conditions and Principal Component Analysis were performed using the package Momocs (43) in Rstudio version 1.0.153.

## Declarations

### Ethics approval and consent to participate

Specimens from Brazil were collected under SISBIO permit # 43105–1. Specimens from other locations did not request any official authorizations.

### Consent for publication

Not applicable.

### Availability of data and material

The data collected and resources used to perform analysis will be available on Dryad Digital Depository.

### Competing interests

Abderrahman Khila is an Associate Editor at BMC Evolutionary Biology.

### Funding

This work was supported by ERC-CoG # 616346, CNPq-PVE # 400751/2014–3 and by CEBA:ANR-10-LABX-25–01 to A. Khila.

### Authors’ contributions

Conception and Design of experiment: A.J.J.C. and A.K.

Acquisition of data: A.J.J.C., D.A., A.V.L., M.K., F.F.F.M. and A.K.

Analysis and Interpretation: A.J.J.C. and A.K.

Drafting manuscript: A.J.J.C. and A.K.

Revising manuscript: A.J.J.C., D.A., A.V.L., M.K., F.F.F.M. and A.K.

## Acknowledgements

We thank A. Herrel, N. Nadeau, J. Abbott and L. Rowe for helpful discussions; A. Le Bouquin, S. Viala, C. Finet, F. Bonneton, A. Decaras, R. Arbore, W. Toubiana for comments on the manuscript; S. Viala for help with scanning electron microscopy and Centre Technologiques des Microstructure at Universite Claude Bernard Lyon 1 for access to SEM microscope, M. Bouchet for help with CT Scan, L. Souquet for help with shape analysis, and A. Le Bouquin and P. Joncour for help with statistics.

## References

1. Arnqvist G, Rowe L. Sexual conflict. Princeton, N.J.: Princeton University Press; 2005. xii, 330 p p.

2. Arnqvist G, Rowe L. Antagonistic coevolution between the sexes in a group of insects. Nature. 2002;415(6873):787–9.

3. Pennell TM, Morrow EH. Two sexes, one genome: the evolutionary dynamics of intralocus sexual conflict. Ecology and evolution. 2013;3(6):1819–34.

4. Bonduriansky R, Chenoweth SF. Intralocus sexual conflict. Trends in ecology & evolution. 2009;24(5):280–8.

5. Rowe L, Day T. Detecting sexual conflict and sexually antagonistic coevolution. Philosophical transactions of the Royal Society of London Series B, Biological sciences. 2006;361(1466):277–85.

6. Arnqvist G, Rowe L. Correlated evolution of male and female morphologles in water striders. Evolution; international journal of organic evolution. 2002;56(5):936–47.

7. Weigensberg I, Fairbairn DJ. The sexual arms race and phenotypic correlates of mating success in the waterstrider, Aquarius remigis (Hemiptera: Gerridae). Journal of Insect Behavior. 1996;9(2):307–19.

8. Arnqvist G, Rowe L. Sexual Conflict and Arms Races between the Sexes - a Morphological Adaptation for Control of Mating in a Female Insect. Proceedings of the Royal Society B-Biological Sciences. 1995;261(1360):123–7.

9. Dean R, Perry JC, Pizzari T, Mank JE, Wigby S. Experimental evolution of a novel sexually antagonistic allele. PLoS genetics. 2012;8(8):e1002917.

10. Mank JE. The transcriptional architecture of phenotypic dimorphism. Nature ecology & evolution. 2017;1(1):6.

11. Mank JE, Wedell N, Hosken DJ. Polyandry and sex-specific gene expression. Philosophical transactions of the Royal Society of London Series B, Biological sciences. 2013;368(1613):20120047.

12. Snook RR, Bacigalupe LD, Moore AJ. The quantitative genetics and coevolution of male and female reproductive traits. Evolution; international journal of organic evolution. 2010;64(7):1926–34.

13. Bergsten J, Miller KB. Phylogeny of Diving Beetles Reveals a Coevolutionary Arms Race between the Sexes. Plos One. 2007;2(6).

14. Kuntner M, Coddington JA, Schneider JM. Intersexual arms race? Genital coevolution in nephilid spiders (Araneae, Nephilidae). Evolution; international journal of organic evolution. 2009;63(6):1451–63.

15. Arnqvist G. SPATIAL VARIATION IN SELECTIVE REGIMES: SEXUAL SELECTION IN THE WATER STRIDER, GERRIS ODONTOGASTER. Evolution; international journal of organic evolution. 1992;46(4):914–29.

16. Andersen NM. The semiaquatic bugs (Hemiptera: Gerromorpha). Klampenborg, Denmark.: Scandinavian Science Press LTD.; 1982.

17. Moreira FFF, Ribeiro JRI. Two new Rhagovelia (Heteroptera: Veliidae) and new records for twelve species in southeastern Brazil. Aquatic Insects. 2009;31(1):45–61.

18. Polhemus DA. Systematics of the genus Rhagovelia Mayr (Heteroptera: Veliidae) in the Western Hemisphere (exclusive of the angustipes complex). Lanham, Md.: Entomological Society of America; 1997. ii, 386 p. p.

19. Andersson MB. Sexual selection. Princeton, N.J.: Princeton University Press; 1994. xx, 599 p. p.

20. Khila A, Abouheif E, Rowe L. Function, developmental genetics, and fitness consequences of a sexually antagonistic trait. Science (New York, N Y). 2012;336(6081):585–9.

21. Ivy TM, Sakaluk SK. Sequential mate choice in decorated crickets: females use a fixed internal threshold in pre- and postcopulatory choice. Animal Behaviour. 2007;74:1065–72.

22. Karlsson B, Johansson A. Seasonal polyphenism and developmental trade-offs between flight ability and egg laying in a pierid butterfly. Proceedings Biological sciences. 2008;275(1647):2131–6.

23. Simmons LW, Emlen DJ. Evolutionary trade-off between weapons and testes. Proceedings of the National Academy of Sciences of the United States of America. 2006; 103(44): 16346–51.

24. Langellotto GA, Denno RF, Ott JR. A trade - off between flight capability and reproduction in males of a wing - dimorphic insect. Ecology. 2000;81(3):865–75.

25. Braendle C, Davis GK, Brisson JA, Stern DL. Wing dimorphism in aphids. Heredity. 2006;97(3):192–9.

26. Kawada K. Forms and morphs of aphids. In: Minks AKH, P., editor. Aphids, Their Biology, Natural Enemies and Control. Vol 2A. Amsterdam: Elsevier; 1987. p. 255–66.

27. Miyazaki M. Forms and morphs of aphids. In: Minks AKH, P., editor. Aphids, Their Biology, Natural Enemies and Control. Vol 2A. Elsevier: Amsterdam ed1987. p. 163–95.

28. Denno RF, Olmstead KL, Mccloud ES. Reproductive Cost of Flight Capability - a Comparison of Life-History Traits in Wing Dimorphic Planthoppers. Ecological Entomology. 1989;14(1):31–44.

29. Moran NA. The Evolutionary Maintenance of Alternative Phenotypes. American Naturalist. 1992;139(5):971–89.

30. Chesebro J, Hrycaj S, Mahfooz N, Popadic A. Diverging functions of Scr between embryonic and post-embryonic development in a hemimetabolous insect, Oncopeltus fasciatus. Developmental biology. 2009;329(1):142–51.

31. Markow TA, Bustoz D, Pitnick S. Sexual selection and a secondary sexual character in two Drosophila species. Animal Behaviour. 1996;52:759–66.

32. Polak M, Starmer WT, Wolf LL. Sexual selection for size and symmetry in a diversifying secondary sexual character in Drosophila bipectinata duda (Diptera : Drosophilidae). Evolution. 2004;58(3):597–607.

33. Spieth HT. Mating Behavior within the Genus Drosophila (Diptera). Bulletin of the American Museum of Natural History. 1952;99(7):401–74.

34. Tanaka K, Barmina O, Sanders LE, Arbeitman MN, Kopp A. Evolution of Sex-Specific Traits through Changes in HOX-Dependent doublesex Expression. Plos Biology. 2011;9(8).

35. Fairbairn DJ, King E. Why do Californian striders fly? Journal of evolutionary biology. 2009;22(1):36–49.

36. Spence JR. The Habitat Templet and Life-History Strategies of Pond Skaters (Heteroptera, Gerridae) - Reproductive Potential, Phenology, and Wing Dimorphism. Canadian Journal of Zoology-Revue Canadienne De Zoologie. 1989;67(10):2432–47.

37. Ditrich T, Papacek M. Effect of population density on the development of Mesovelia furcata (Mesoveliidae), Microvelia reticulata and Velia caprai (Veliidae) (Heteroptera: Gerromorpha). European Journal of Entomology. 2010;107(4):579–87.

38. Venables WN, Ripley BD. Modern applied statistics with S. 4th ed. New York: Springer; 2002. xi, 495 p. p.

39. Crumiere AJJ, Santos ME, Semon M, Armisen D, Moreira FFF, Khila A. Diversity in Morphology and Locomotory Behavior Is Associated with Niche Expansion in the Semi-aquatic Bugs. Current biology : CB. 2016;26(24):3336–42.

40. Maddison WP, Maddison DR. Mesquite: a modular system for evolutionary analysis. Version 3.40. http://mesquiteproiect.org 2018 {

41. Revell LJ. phytools: an R package for phylogenetic comparative biology (and other things). Methods in Ecology and Evolution. 2012;3(2):217–23.

42. Metscher BD. MicroCT for comparative morphology: simple staining methods allow high-contrast 3D imaging of diverse non-mineralized animal tissues. BMC physiology. 2009;9:11.

43. Bonhomme V, Picq S, Gaucherel C, Claude J. Momocs: Outline Analysis Using R. Journal of Statistical Software. 2014;56(13):1–24.

